# Reduced *PIN1* gene expression in neocortical and limbic brain regions in female Alzheimer’s patients correlates with cognitive and neuropathological phenotypes

**DOI:** 10.1101/2023.08.14.553279

**Authors:** Camila de Ávila, Crystal Suazo, Jennifer Nolz, J. Nicholas Cochran, Qi Wang, Ramon Velazquez, Eric Dammer, Benjamin Readhead, Diego Mastroeni

## Abstract

Women have a higher incidence of Alzheimer’s disease (AD), even after adjusting for increased longevity. Thus, there is an urgent need to identify the molecular networks that underpin the sex-associated risk of AD. Recent efforts have identified *PIN1* as a key regulator of tau phosphorylation signaling pathway*. Pin1* is the only gene, to date, that when deleted can cause both tau and Aβ-related pathologies in an age-dependent manner. We analyzed multiple brain transcriptomic datasets focusing on sex differences in *PIN1* mRNA levels, in an aging and AD cohort, which revealed reduced *PIN1* levels driven by females. Then, we validated this observation in an independent dataset (ROS/MAP) which also revealed that *PIN1* is negatively correlated with multiregional neurofibrillary tangle density and global cognitive function, in females only. Additional analysis revealed a decrease in *PIN1* in subjects with mild cognitive impairment (MCI) compared with aged individuals, again, driven predominantly by female subjects. Our results show that while both male and female AD patients show decreased *PIN1* expression, changes occur before the onset of clinical symptoms of AD in females and correlate to early events associated with AD risk (e.g., synaptic dysfunction). These changes are specific to neurons, and may be a potential prognostic marker to assess AD risk in the aging population and even more so in AD females with increased risk of AD.

**Graphical abstract:** 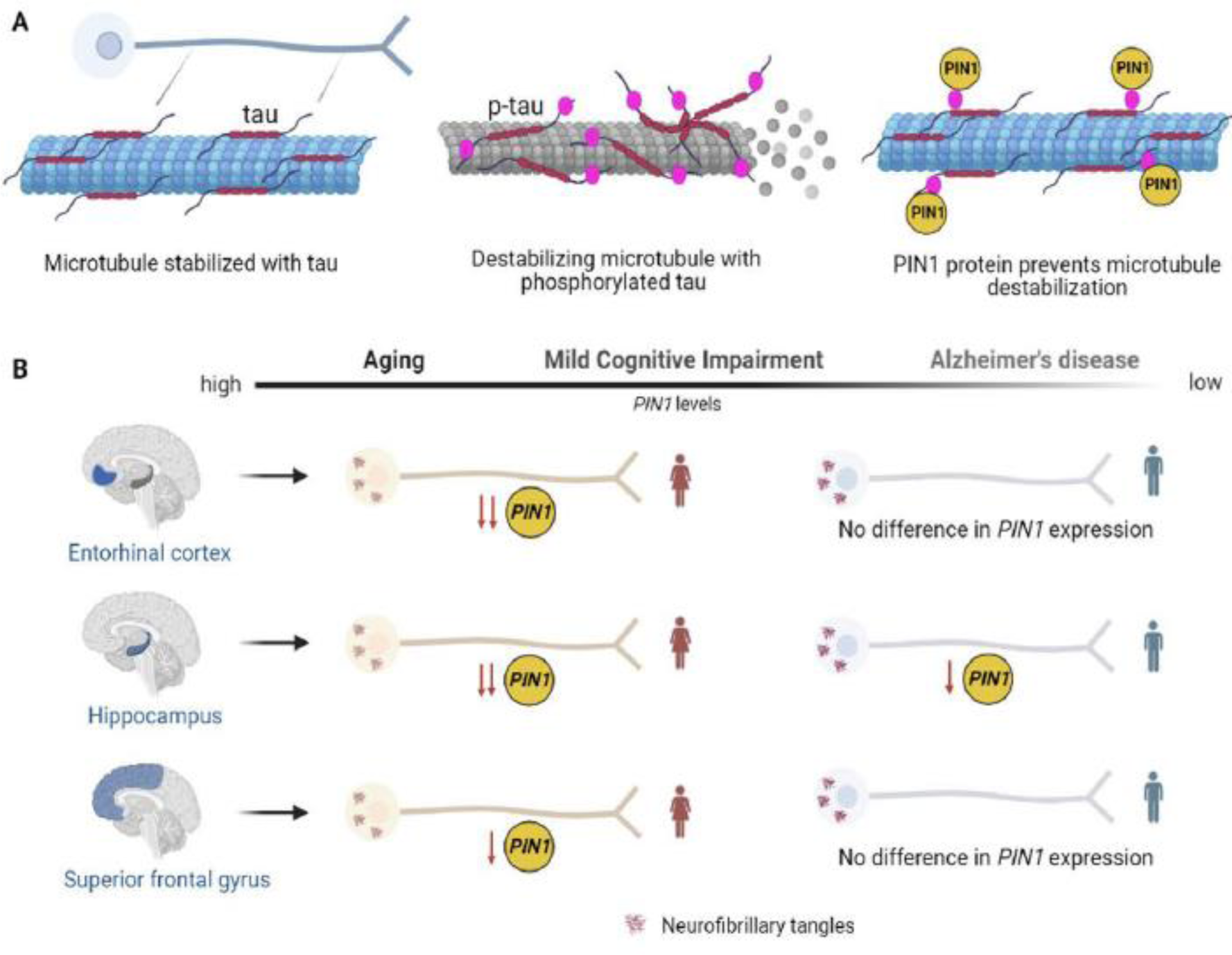

## 1. INTRODUCTION

Alzheimer’s disease (AD), the most prevalent neurodegenerative disorder worldwide, is clinically characterized by impairments in cognition, memory, and intellectual abilities [1]. The neuropathological hallmarks of AD are brain extracellular amyloid-β peptide (Aβ) plaques, intraneuronal tangles comprised of hyperphosphorylated tau, and synaptic and neuronal loss [1]. Over the next few decades, the advancing age of the global population will drive a dramatic increase in AD burden if effective interventions remain limited; it is estimated that by the year 2050, the prevalence within the US will rise to 20 million individuals [2]. This is alarming given that current treatments have limited ability to prevent, treat, or manage AD.

Importantly, sex-differences have been reported in different aspects of AD such as epidemiology, risk factors progression, symptomatology, and biomarkers [3]. Women have a higher incidence [4] and prevalence of Alzheimer’s dementia than men, even after adjustment for increased longevity [3, 5, 6, 7]. Thus, there is an unmet and urgent need to identify the neural networks that govern cognitive processes and contribute to the sex-related risk of AD.

Moreover, phosphorylation of amyloid precursor protein’s variants (APP) and hyperphosphorylation of tau protein are both increased in AD brains, can lead to formation of Aβ and neurofibrillary tangles (NFTs) respectively [8].These observations suggest that phosphorylation events are critical to the understanding of the pathogenesis and treatment of AD. One critical protein involved in phosphorylation pathways is the peptidyl-prolyl *cis/trans*isomerase (PIN1), which catalyzes the isomerization of the peptide binding bond between phosphorylated Ser/Thr-Pro in proteins, thereby regulating the function of proteins after phosphorylation [9]. Various reports have found that Pin1 expression is dysregulated in AD [10, 11, 12]. Indeed, Pin1 expression levels are reduced in AD hippocampus [10].

*Pin1* is the only gene, to date, that when deleted can cause both tau and Aβ-related pathologies in an age-dependent manner, in mice [9, 13]. Additionally, *Pin1* homozygous knockout (KO) mice develop a progressive phenotype of premature aging [14]. *Pin1* KO mice show age-dependent neuropathy characterized by tau hyper-phosphorylation, tau filament formation, APP amyloidogenesis, intracellular Aβ_42_ accumulation, and neuronal degeneration [9, 13, 15]. Notably, Pin1 has been shown to play a protective role for neurons under toxic *in vitro* conditions [10, 14]. Collectively, these findings suggest that alterations in *PIN1* expression may be one of the driving forces in the initiation and progression of AD pathogenesis.

Although previous work has shown that PIN1 expression levels are significantly reduced in AD human post-mortem brain tissue [16, 17], it has yet to be determined whether *PIN1* expression is dysregulated as a function of age and sex. Additionally, it is not known which specific classes of central nervous system cells show reduced expression. Thus, the goal of this study was to first determine whether the expression of *PIN1* is significantly altered in aging (20-90 years). Additionally, we were interested in determining whether the expression of *PIN1* was altered in pre-AD cases (amnestic MCI). Finally, we validated AD expression changes using three independent studies using multiple expression platforms. The results from this comprehensive study indicate that *PIN1* may be an informative molecule for illuminating pre-clinical AD in females, as well as a marker of disease progression.

## 2. METHODS

### 2.1 Tissue Homogenates and Affymetrix arrays

#### Tissue Collection

As previously published [18, 19], frozen unfixed tissue was available from two or more regions from 85% of the cases, resulting in a total of 193 tissue samples (56 EC, 62 HIPP, and 75 SFG). SFG (Superior Frontal Gyrus, crest/superior surface at the genu of the corpus callosum), HIPP (Hippocampus, body of the HIPP at the level of the lateral geniculate nucleus), EC (Entorhinal Cortex, crest of the parahippocampal gyrus at the level of the anterior hippocampus). Group sizes were as follows: young (n = 61, 20 to 59 years, mean age 35.4 ± 10.5 years), aged controls (n = 73, age 69 to 99, mean age 84.2 ± 8.9 years), and AD cases (n = 59, ages 74 to 95 years, mean age 85.7 ± 6.5 years) with males and females similarly represented in each group. Refer to references [18, 19] for detailed sample information. Total RNA was extracted from the hippocampus as described previously^19^.

All genes that did not meet the 50% present call threshold were removed by Genespring G 7.3.1 Expression Analysis software. We preferentially selected those probe sets that were most specific, such that they were annotated with the smallest number of Ensembl gene IDs. After applying this criterion, if there remained multiple probes for any one gene, we excluded those probe sets expected to hybridize with targets in a non-specific fashion (i.e., those with “_x_” in the Affymetrix identifier). If there remained multiple probes for a given gene, we took the mean of the probe sets.

#### Statistical analysis

Select genes were investigated for statistical significance (*p* < 0.01). A two-tailed paired t-test, assuming equal variance (using multiple testing corrections, by Benjamini and Hochberg False Discovery Rate), was applied to locate genes that were significant in differentiating expression between young and aged controls and AD and aged controls.

Data have been deposited in the Gene Expression Omnibus database (www.ncbi.nlm.nih.gov/geo) accession number GSE11882.

### 2.2. Laser capture of glial and neuronal cells and RNA sequencing

Frozen hippocampal tissue sections (10µm) were mounted onto PEN slides and fixed in ice-cold acetone/ethanol for 10 min on ice. Sections were washed in ice-cold 1X Phosphate buffered saline (PBS), blocked in 1% hydrogen peroxide for 2 minutes, followed by 3 quick submersions in ice-cold 1X PBS. Sections were then placed in 1:500 dilution of primary antibody in 1X PBS supplemented with 50U RNAseOUT (Invitrogen) for 10 minutes at room temperature. After the incubation, sections were washed three times in 1X PBS and incubated in avidin (1:300)-biotin (1:300) complex (Vector) in 1X PBS for 10 minutes at room temperature (RT). Sections were washed three times in 50mM Tris buffer and immersed in 3.3’-diaminobenzidine (DAB) solution (9.3 ml 50mM Tris; 200μl DAB (5mg/ml); 500μl saturated nickel; and 4 μl of 1% H_2_0_2_) for 5 minutes, followed by two quick rinses in 50mM Tris to stop the reaction.

#### Laser Capture of LN3-positive microglia

LN3 antibody reacts with MHCII microglia. These microglia are said to be antigen-presenting (e.g., activated, M1 phase, but not phagocytic M2). Immediately after the detection of LN3-immunoreactivity, sections were dipped in 100% ethanol and loaded onto a Leica AS-LMD laser capture microscope. After objective calibration, 600 LN3-positive cells were captured using 20X magnification. Individual microglial cells were cut and dropped into an inverted microcentrifuge cap containing 50μl buffer RLT (RNeasy Micro Kit - Qiagen) and 1% β-Mercaptoethanol.

#### Laser Capture of GFAP-positive Astrocytes

Glial fibrillary acidic protein is an intermediate filament protein that is expressed by astrocytes. Immediately after the detection of GFAP-immunoreactivity, sections were dipped in 100% ethanol and loaded onto a Leica AS-LMD laser capture microscope. After objective calibration, 600 GFAP-positive cells were captured using 20X magnification. Individual microglial cells were cut and dropped into an inverted microcentrifuge cap containing 50 μl buffer RLT (RNeasy Micro Kit - Qiagen) and 1% β-Mercaptoethanol.

#### Laser Capture of neurons

Brain sections were stained with 1% neutral red (Fisher Scientific); pyramidal neurons were identified by their characteristic size, shape, and location. Immediately after staining sections were dipped in 100% ethanol and loaded onto a Leica AS-LMD laser capture microscope. After objective calibration, 300 pyramidal neurons were captured using 20X magnification. Individual microglial cells were cut and dropped into an inverted microcentrifuge cap containing 50 μl buffer RLT (RNeasy Micro Kit - Qiagen) and 1% β-Mercaptoethanol.

#### Laser Captured RNA sequencing

RNAseq library preparation, paired end sequencing, quality control procedures, gene expression quantification and differential expression analysis were performed as previously described [20].

### 2.3. Laser Capture of neurons containing neurofibrillary tangles and neurons without neurofibrillary tangles and Affymetrix arrays

#### Tissue Collection

As previously published [21, 22], brain samples were collected at three Alzheimer’s disease centers (Washington University, Duke University, and Sun Health Research Institute) from clinically and neuropathologically classified late-onset AD-afflicted individuals and well-matched non-demented (ND) subjects. Detailed information on controls and AD subjects can be found in previous reports [21, 22, 23].

As previously described [21, 22], brain sections were stained with a combination of thioflavin-S (Sigma) and 1% neutral red (Fisher Scientific); pyramidal neurons were identified by their characteristic size, shape, and location within the region of interest, and tangles were identified by the green fluorescence of thioflavin-S staining. In the HIPP, the pyramidal cells lacking thioflavin-S staining, and neurons with thioflavin-S staining, were collected. For each individual, 500 pyramidal neurons were collected using laser capture microdissection (Veritas automated laser capture microdissection system; Arcturus).

#### Expression Profiling

Expression profiling was performed as previously described [21, 22]. Isolated total RNA was double-round amplified, cleaned, and biotin-labeled with the Affymetrix GeneChip two-cycle target labeling kit with a T7 promoter and the Ambion MEGAscript T7 high-yield transcription kit according to the manufacturer’s instructions. Amplified and labeled cRNA was quantitated on a spectrophotometer and run on a 1% TAE gel to check for an evenly distributed range of transcript sizes, for detailed case information see Liang et al. 2008. Twenty μg of cRNA were fragmented to ≈35–200 bp by alkaline treatment (200 mM Tris-acetate, pH 8.2; 500 mM KOAc; 150 mM MgOAc) and run on a 1% TAE gel to verify fragmentation. Separate hybridization cocktails were made by using 15 μg of fragmented cRNA from each sample according to Affymetrix’s instructions.

#### Microarray Analysis

Briefly, 200μl of each mixture was separately hybridized to an Affymetrix Human Genome U133 Plus 2.0 array for 16 h at 45°C in the Hybridization Oven 640. Arrays were washed on the Affymetrix upgraded GeneChip Fluidics Station 450 by using a primary streptavidin phycoerythrin (SAPE) stain, subsequent biotinylated antibody stain, and secondary SAPE stain. Arrays were scanned on the Affymetrix GeneChip Scanner 3000 7G with AutoLoader. For more details refer to previous publications [21, 22, 24].

#### Statistical Analysis

Direct comparisons between neurologically healthy and AD-afflicted brains were performed in the hippocampus to analyze expression differences. For each analysis, genes that did not demonstrate at least ≈40% present calls across all transcripts profiled for each region-specific comparison were removed by using Genespring GX 7.3 Expression Analysis software (Agilent Technologies). A two-tailed unpaired T-test, assuming unequal variances (with a multiple testing correction using the Benjamini and Hochberg FDR), was applied to each comparison for all genes that passed the 40% present-call criterion to locate genes [25, 26] that were statistically significant in differentiating expression between healthy brains and AD brains. (After the present-call filter, 31,496 genes were identified in the hippocampus). For each analysis comparing AD expression levels with control levels, genes that had a corrected *P*-value ≤ 0.01 were collected, and those genes whose average AD signal and average control signal were both below a threshold of 150 were removed.

Microarray gene expression data downloaded from Gene Expression Omnibus (Preclinical AD: accession GSE9770) [21, 22], Control accession: GSE5281 [21, 22].

### 2.4. Immunohistochemical studies

Chromogenic immunohistochemical studies were completed on twenty-four human temporal neocortical samples, 12 AD and 12 ND samples. Samples were matched for PMI, age, and sex, among other covariates (see table of samples, **Supplementary Table 1)**. For detailed methods please see references [27, 28, 29]. Briefly, 40µm free-floating sections were blocked in H_2_O_2_ and bovine serum albumin. Following blocking steps, tissues were incubated in antibody raised against Pin1 (1:200 dilution, Sigma), overnight (ON) at 4°C. Sections were washed and then incubated in species-specific secondary (1:1000, Vector) for two hours at room temperature. Sections were washed and incubated in 1:1000 avidin/biotin reagent, washed and incubated in DAB. All sections were reacted for the same amount of time, dried, taken through graded alcohols, cleared in xylene, and mounted using permount. Adjacent serial sections were stained with cresyl violet, or within sections with neutral red for structural visualization. Slides were imaged using an Olympus IX71.

<u>For fluorescence microscopy</u>, the sections were washed 3X in PBST, blocked with either 3% normal goat serum or 3% BSA, and incubated for 1h. After further washing, sections were incubated in primary antibody [*Pin1*(mse) antibody, 1:200 dilution, PS396 (rb) 1:1000, T231 (rb) 1:500] ON at 4°C. Sections were washed 3X in PBST and incubated in species-specific, fluorophore-conjugated secondary antibodies (Molecular Probes). After a final wash, the sections were mounted, taken through Sudan Black to reduce autofluorescence, and coverslipped with Vectashield (Vector). Immunostained tissue sections were examined on Nikon Eclipse Ti2 confocal and Olympus IX70 microscopes equipped with epifluorescence illumination. The findings were documented photographically with an Olympus DP-71 color digital camera or, for confocal microscopy, by Nikon A1/A1R. Blocking peptides were used to confirm antibody specificity (**Supplementary** Figure 1)

### 2.5. Western blot analysis

For Western blots, frozen temporal cortical blocks were lysed in a solution containing 20mM Tris pH7.5, 0.5% Nonident (Sigma), 1mM EDTA (Sigma), 0.1M NaCl (Sigma), 1mM PMSF (Sigma), Sigma protease inhibitors 1, 2, and complete protease inhibitor cocktail (Roche). Protein concentrations were determined by BCA assay (Pierce) using bovine serum albumin as the standard. A total of 40µg of sample protein was combined with Laemmli sample buffer for separation by SDS-PAGE, followed by transfer to PVDF membrane (Bio-Rad). Membranes were blocked using 5% non-fat dry milk and probed with anti-Pin1. After incubation with the primary antibody, membranes were washed, incubated with the secondary antibody, washed again, reacted with chemiluminescence substrate (Pierce), and imaged on Amersham imager 680 (GE). Exposure 1.5 minutes.

### 2.6. *PIN1* expression analysis in ROS/MAP dataset

RNA sequencing data was obtained from the Accelerating Medicines Partnership - Alzheimer’s Disease (AMP-AD) Knowledge Portal (ROS/MAP: syn3505720). ROS and MAP cohort samples were collected from the dorsolateral prefrontal cortex (DLPFC) as previously described [30]. Gene expression within the ROS and MAP cohorts was available as gene FPKM estimates. We log2 transformed *PIN1* FPKM values after incrementing an offset of 0.5 to avoid the transformation of zero values. Subjects were classified according to a clinical cognitive diagnosis summary as AC (Aged controls, no cognitive impairment), MCI (mild cognitive impairment, no other condition contributing to cognitive impairment), and AD (Alzheimer’s disease dementia, no other condition contributing to cognitive impairment).

## 3. RESULTS

### 3.1. *PIN1* mRNA levels are downregulated as a function of age

The most salient risk factor for Alzheimer’s disease and other dementias is aging. To understand the effect of aging on the expression of *PIN1,* we analyzed three brain regions: HIPP, ENT, and SFG in an aging cohort: young control (YC) (n = 22, 20 to 59 years, mean age 35.4 ± 10.5 years) vs. aged control (AC) (n = 33, age 69 to 99, mean age 84.2 ± 8.9 years)[25, 31]. Significant downregulation of *PIN1* was observed in AC compared to YC males and females in the HIPP, and SFG. Only females were significantly downregulated in ENT (**Fig. 1A**). Linear regression analysis (Pearson’s correlation coefficient (r)) revealed a significant correlation with *PIN1* as a function of age in HIPP (P<0.001, F-ratio 8.48) (**Fig. 1B**), ENT (P<0.0001, F-ratio 22.65) (**Fig. 1C**), and SFG (P<0.001, F-ratio 11.20) (**Fig. 1D**). These data show that *PIN1* mRNA levels significantly decrease as a function of age.

**Figure 1.**
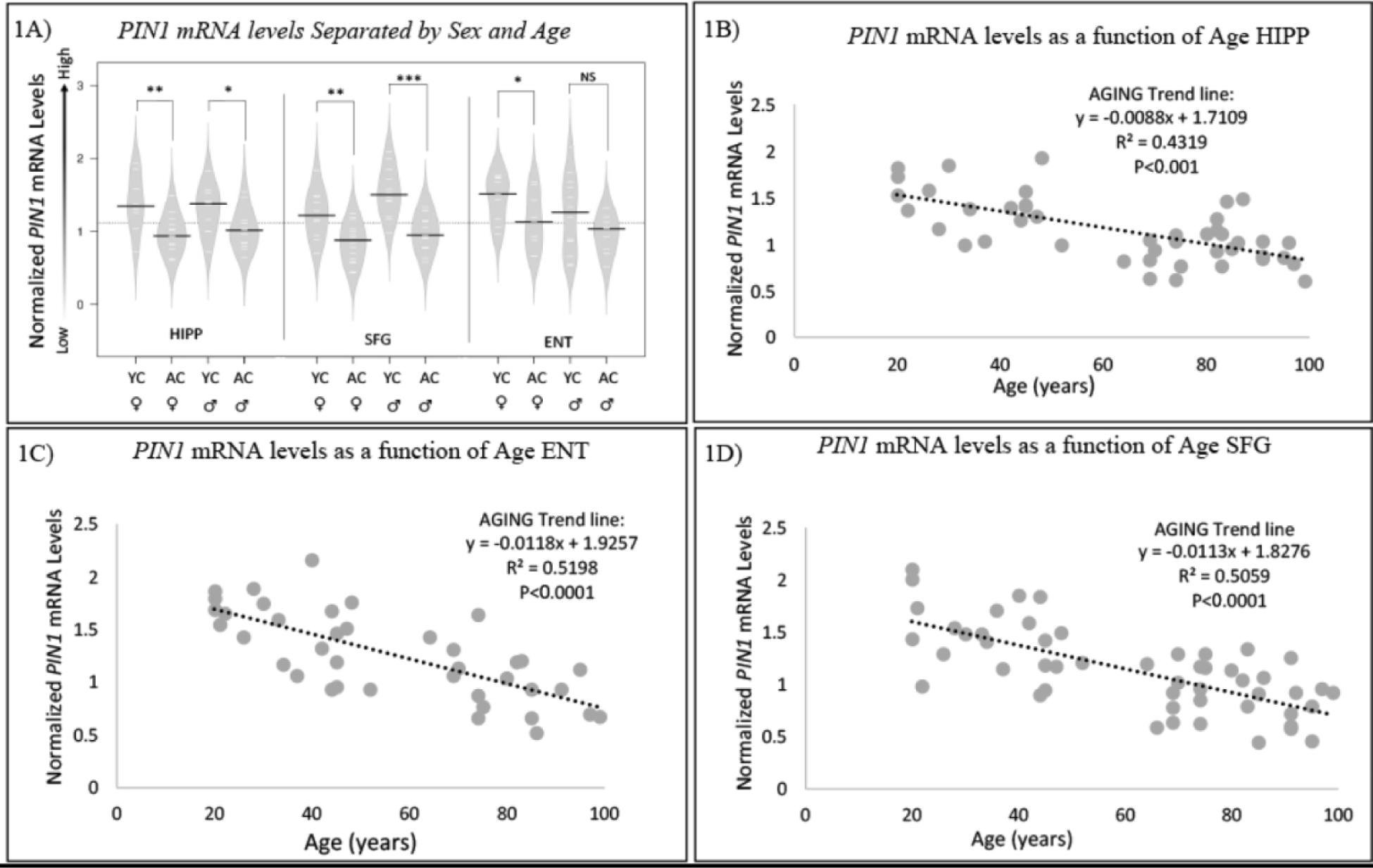
*PIN1* mRNA levels are downregulated as a function of age. To understand the effect of age and sex we analyzed three brain regions: hippocampus (HIPP), entorhinal cortex (ENT), and superior frontal gyrus (SFG) in an aging cohort: young (n = 22, 20 to 59 years, mean age 35.4 ± 10.5 years) vs. aged controls (n = 33, age 69 to 99, mean age 84.2 ± 8.9 years). 1A) Black lines show group medians; white lines represent individual data points; polygons represent the estimated density of the data; dotted horizontal line represents population mean normalized mRNA abundance. Mean expression levels within groups revealed a significant downregulation of *PIN1* as a function of age in AC females and males. Female subjects (YC and AC) showed lower *PIN1* expression levels compared to their male counterparts in all regions except for ENT (1A). Females, however, did show a significant difference (1A). Linear regression analysis revealed a significant negative correlation of *PIN1* with age in HIPP (B), ENT (C), and SFG (D) regardless of sex. (p<0.01*, p<0.001**, p<0.0001 ***, NS= not significant).

### 3.2. *PIN1* mRNA levels are reduced in Alzheimer’s disease compared to age-matched controls

To characterize the effect of AD on *PIN1* expression, we compared the same aged controls (AC, n = 33, age 69 to 99, mean age 84.2 ± 8.9 years), to clinically and neuropathologically confirmed AD cases (n = 26, ages 74 to 95 years, mean age 85.7 ± 6.5 years). As observed for advanced age (**Fig.1**), *PIN1* levels were significantly down-regulated in AD-hippocampus in AD vs. AC (p<0.0005, Log 2-Fold Change -2.4). When separated by sex, both females (p<0.0001, F-ratio 11.79) and males (p<0.049, F-ratio 4.94) were significantly downregulated, but females were overall the most affected (**Fig. 2**). In the SFG, AD vs. ND showed significant *PIN1* downregulation (p<0.01, F-ratio 6.32) overall. When separated by sex, only females were significantly downregulated (p<0.01, F-ratio 8.12) (**Fig.2**). Similarly, In the ENT, AD vs. ND showed an overall significant difference (p<0.001, F-ratio 10.19). When separated by sex, only females were significantly downregulated (p<0.001, F-ratio 8.42) (**Fig. 2**). These findings indicate that *PIN1* expression changes are largely driven by females in the HIPP and are solely responsible for the overall AD-associated expression changes in the SFG and ENT.

**Figure 2.**
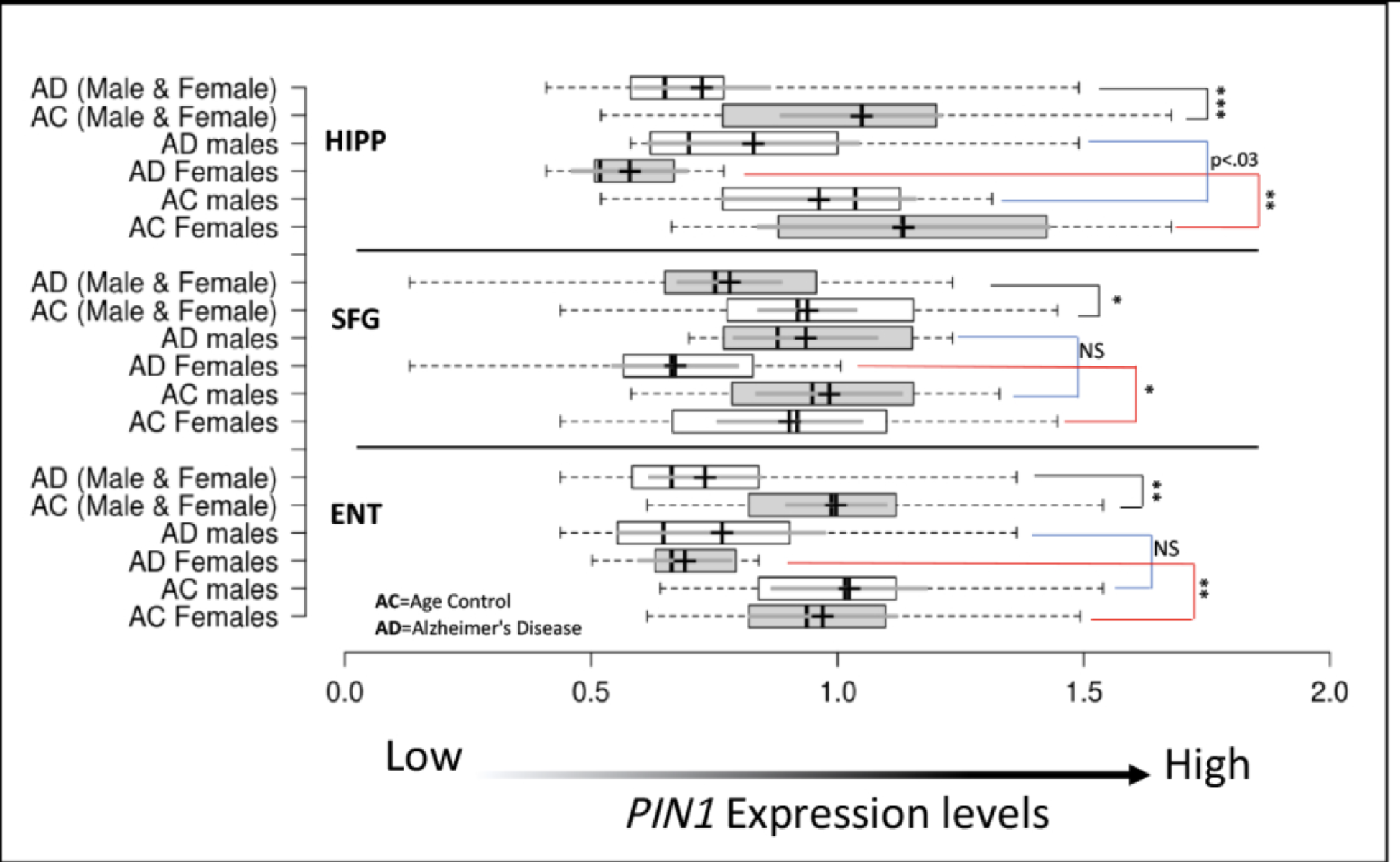
*PIN1* mRNA levels are reduced in Alzheimer’s disease compared to age-matched controls. To address the effect that AD has on the expression of *PIN1*, we compared the same aged controls (n = 33, age 69 to 99, mean age 84.2 ± 8.9 years), to clinically and neuropathologically confirmed AD cases (n = 26, ages 74 to 95 years, mean age 85.7 ± 6.5 years). Center lines show the medians; box limits indicate the 25th and 75th percentiles; whiskers extend to minimum and maximum values; crosses represent sample means; bars indicate 95% confidence intervals of the means. Data show that females drive the overall significant changes in HIPP, SFG and ENT. The only significant male-associated change was observed in the HIPP (p=0.03). Black bars are a comparison between AD and AC, these include both males and females. The blue bars are a comparison between AD males and AC males. The red bars are a comparison between AD females and AC females. *=p<0.01, **=p<0.001, ***=p<0.0001, NS= Not Significant.

### 3.3. Impaired cognition and increased neuropathology severity are associated with reduced *PIN1* expression in females

To further explore how the expression of *PIN1* varies with age, sex, and clinicopathological markers of AD, we incorporated post-mortem brain RNA-sequence data from the Religious Orders Study^47^ (ROS), and Memory and Aging Project [30, 32] (MAP) cohorts. These data together (ROS/MAP) comprised 638 brain tissue samples collected from the DLPFC. We obtained normalized gene expression data and examined the abundance of *PIN1* in samples obtained from subjects with varying diagnoses, severity of neuropathology, sex, and cognitive scores.

We observed a nominal reduction of *PIN1* expression in subjects with MCI compared with non-cognitively impaired subjects (MCI vs. AC, T-statistic: -2.21, P-value: 0.027), and a more dramatic decrease in *PIN1* among subjects diagnosed with AD (AD vs. AC, T-statistic: -3.67, P-value: 3e-4) (**Fig. 3A**). When we stratified *PIN1* expression by sex, we observed that the reduction in *PIN1* expression in AD vs. AC was driven entirely by female subjects (**Fig. 3B**, Female AD vs. AC, T-statistic: -3.75, P-value: 3e-4; Male AD vs. AC, T-statistic: 0.90, P-value: 0.37).

**Figure 3.**
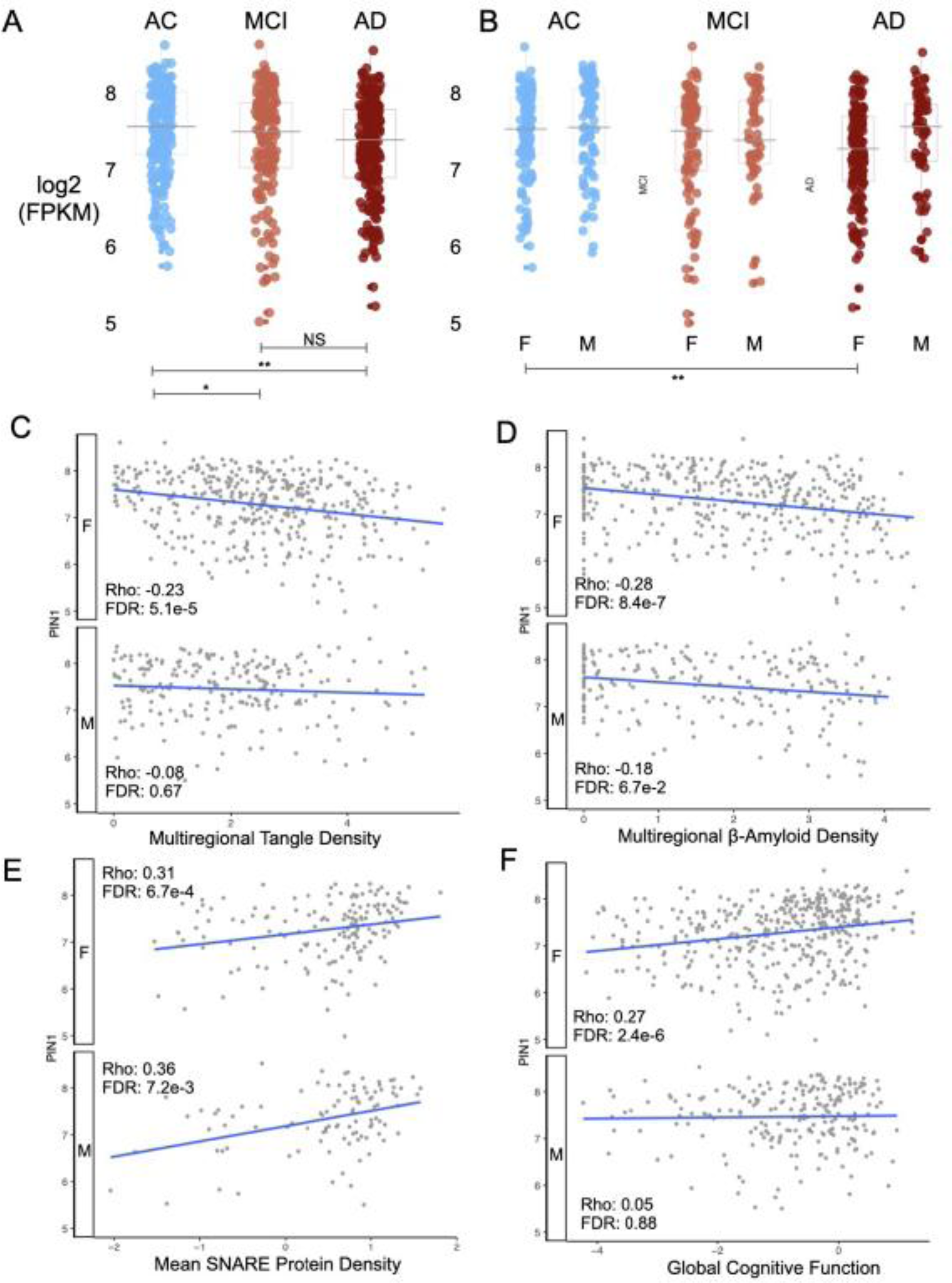
Impaired cognition and increased neuropathology severity are associated with reduced *PIN1* expression in females. (A) *PIN1* expression in DLPFC samples from 638 AD, MCI and AC subjects profiled within the ROS/MAP cohorts, (B) stratified by sex. Sex-stratified spearman correlations between *PIN1* expression and (C) multiregional neurofibrillary tangle density, (D) multiregional β-Amyloid density, (E) mean SNARE protein density, and (F) global cognitive function.

We also observed that *PIN1* expression varied with markers of neuropathology severity in a sex-specific manner. While *PIN1*expression varied inversely with Braak and Braak score (Braak score) across the full data set (Braak score vs. *PIN1* expression correlation, Rho: -0.2, P-value: 3.5e-7), this relationship was driven entirely by the female subjects (Braak score vs. *PIN1* expression correlation, Female-only Rho: -0.23, FDR: 5.1e-5, Male-only Rho: -0.08, FDR: 0.67, **Fig. 3C**). In addition, we observed a significant, inverse relationship between *PIN1* expression and a multiregional summary statistic of β-Amyloid plaque density in both male and female subjects (β-Amyloid plaque density vs. *PIN1* expression correlation, Female-only Rho: -0.28, FDR: 8.4e-7, Male-only Rho: -0.18, FDR: 6.7e-2, **Fig. 3D**).

Given the associations between *PIN1* activity and synaptic function, we examined correlations between *PIN1* and a multi-regional measure of soluble N-ethylmaleimide-sensitive factor attachment protein receptor (SNARE) proteins that have been assayed across the MAP cohort (N=258 subjects). Mean SNARE protein immunodensities are based on the average of syntaxin-1, vesicle-associated membrane protein (VAMP), and synaptosomal-associated protein-25 (SNAP-25), aggregated across six different cortical regions (hippocampus, middle frontal gyrus, inferior frontal gyrus, calcarine cortex, ventromedial caudate, and posterior putamen) and converted to a Z-score across all subjects [33]. Mean SNARE protein density has been demonstrated as an indicator of pan-synaptic function, and predictive of cognitive function before death, with declines in SNARE density correlating with synaptic loss in a manner that appears independent from neuropathology-driven synaptic loss [34]. We observed that *PIN1* expression was positively correlated with mean SNARE protein density Z-score in both males and females (Mean SNARE protein immunodensity Z-score vs. *PIN1* expression correlation, Female-only Rho: 0.31, FDR: 6.7e-4, Male-only Rho: 0.36, FDR: 7.2e-3, **Fig. 3E**).

We also observed that *PIN1* was positively correlated with global cognitive function in a sex-specific manner. Global cognitive function is comprised of a Z-score transformed composite from a battery of cognitive tests [35] and was positively correlated with *PIN1* expression, but only in females (Global Cognitive Function Z-Score vs. *PIN1* expression correlation, Female-only Rho: 0.27, FDR: 2.4e-6, Male-only Rho: 0.05, FDR: 0.88, **Fig. 3F**).

### 3.4. *PIN1* is downregulated in laser capture neurons but not glia in the AD hippocampus

To understand the cell-type context of altered PIN1 expression, we used recently published RNA sequencing data from laser-captured neurons, microglia, and astrocytes from the same subjects [20, 36, 37, 38]. Subsequent RNA sequencing analysis of immunohistochemically defined laser-captured neurons showed similar expression changes observed in hippocampal homogenates, with *PIN1* downregulated (log_2_ fold change -1.8, p<0.0001), but showed virtually undetectable expression levels in microglia or astrocytes (**Fig. 4**). Consistent with our earlier findings (**Figs. 1-3**), female neurons show significantly less *PIN1* expression levels compared to male counterparts. This again suggests that female subjects are the likely basis for the observed population level differences in *PIN1* expression in AD and aging. Whether or not *PIN1* has a regulatory role in microglia or astrocytes is unknown, but these data show the relative expression levels are exceedingly low in glial cells compared to neurons.

**Figure 4.**
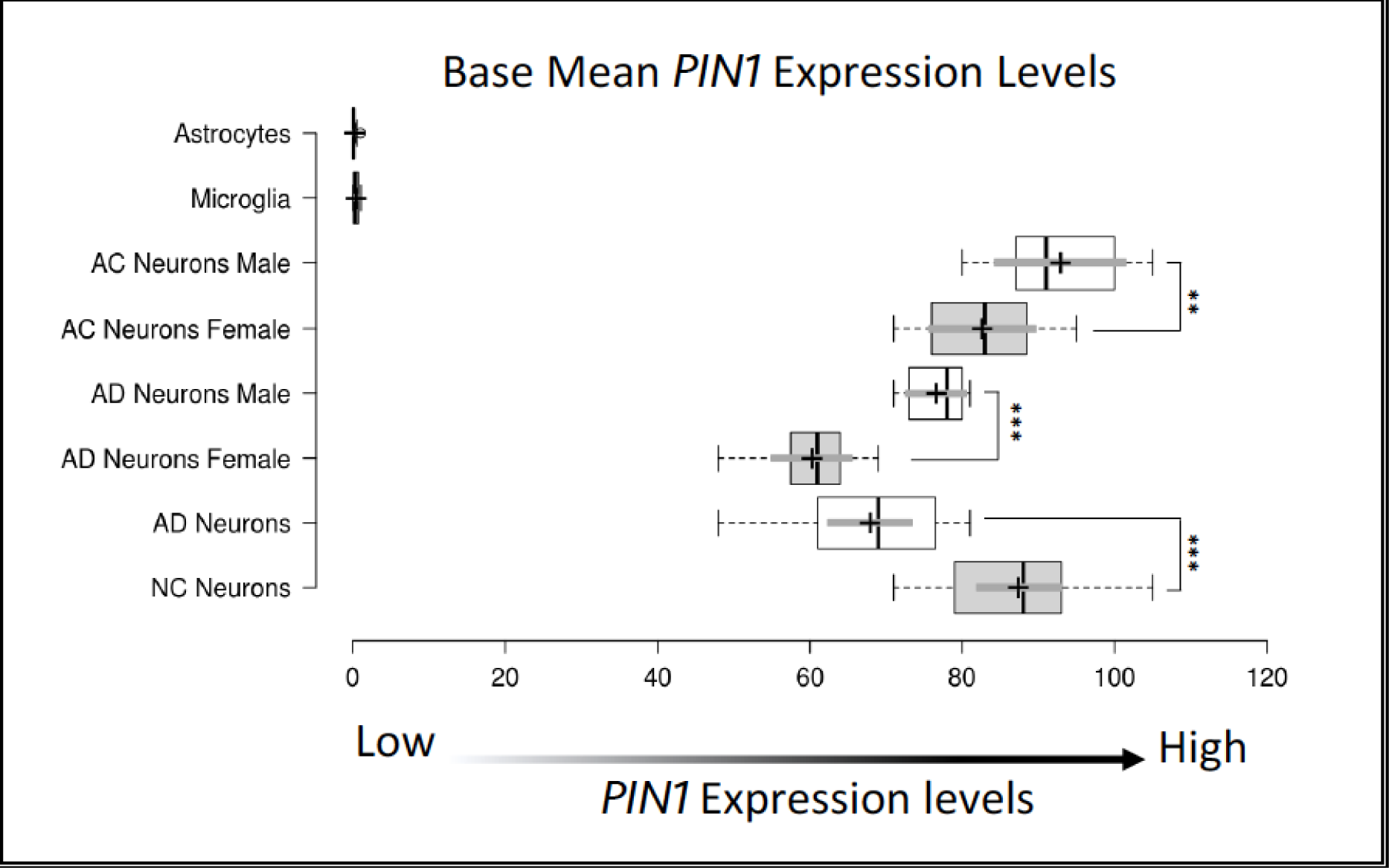
*PIN1* is downregulated in laser capture neurons but not glia in the AD hippocampus. Base mean *PIN1 e*xpression levels per sample in LCM neurons, astrocytes, and microglia. Center lines show the medians; box limits indicate the 25th and 75th percentiles as determined by R software; whiskers extend 1.5 times the interquartile range from the 25th and 75th percentiles; crosses represent sample means; bars indicate 95% confidence intervals of the means. n = 15, 15, 8, 7, 8, 7, 15, 15 sample points. ** p<0.001, ***p<0.0001

### 3.5. *PIN1* expression levels are significantly downregulated in AD neurofibrillary tangle-bearing neurons, but even more so in AD non-tangle neurons

To better understand why AD homogenates and AD laser captured neurons show decreased expression of *PIN1,* we analyzed previously acquired data from our collaborative tangle vs. non-tangle bearing neuron laser capture study [24]. We observed a significant decrease in *PIN1* expression in laser-captured tangle-bearing neurons compared to AC neurons (*p<*0.0001, log_2_ Fold change -2.9, F-ratio 12.34) (**Fig. 5**). Interestingly, we observed the lowest expression levels in non-tangle bearing neurons compared to AC neurons (p<0.0001, log_2_ Fold Change -3.1, F-ratio 12.98). which may indicate that alterations in *PIN1* expression may precede the formation of NFTs. The fact that *PIN1* expression levels are altered in tangle-bearing neurons validates multiple reports on the interaction between *PIN1* and NFTs [14, 39]. No significant differences were observed between male and female tangles, but in non-tangled neurons, females were modestly lower (p<0.033). No significant difference was observed in AC females vs. AC males.

**Figure 5:**
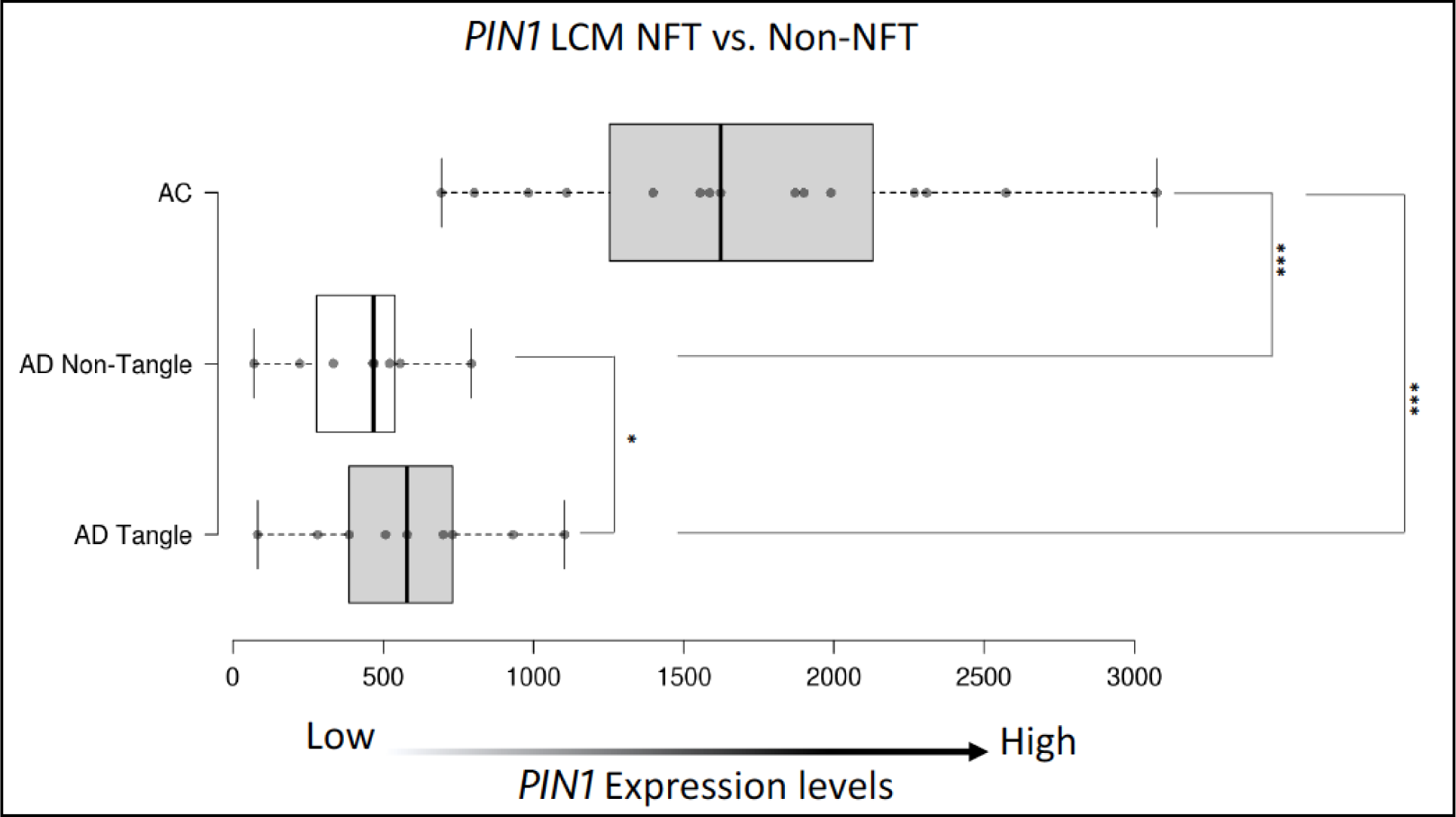
*PIN1* expression levels are significantly downregulated in AD neurofibrillary tangle-bearing neurons, but even more so in AD non-tangle neurons. Laser capture of neurofibrillary tangle (NFT) in AD and non-tangle bearing neurons in AD and AC. Center lines show the medians; box limits indicate the 25th and 75th percentiles; whiskers extend 1.5 times the interquartile range from the 25th and 75th percentiles, outliers are represented by dots; data points are plotted as open circles. n = 9, 8, 15 sample points. AC samples show significantly higher *PIN1* expression levels compared to AD Non-tangle and AD tangle. Interestingly, AD non-tangles showed the lowest *PIN1* expression levels overall. *p<0.01, ***p<0.0001.

### 3.6. Axonal PIN1 protein levels are decreased in females

To investigate sex differences in PIN1 distribution, in AC and AD, we performed immunohistochemistry in the temporal neocortex (**Fig.6**) in 24 AD samples and 24 AC samples (12 females and 12 males/group). PIN1 was compactly wrapped around the nuclear compartment reminiscent of rough endoplasmic reticulum (ER), in AC males and females (red arrow, **6A & 6C** insert). Differences were found in PIN1-Immunoreactivity (IR), specifically in axons; AC-females showed lower levels of PIN1, compared to AC-males (**6A** red arrowheads **vs. 6B**). Interestingly, in AD, PIN1-IR appears to be redistributed outside the ER in no structural organelle in both male and female AD (**Fig. 6C** & **Fig. 6D** insert), like what we have observed in other AD studies [28, 29]. These data may indicate that in normal aging the loss of PIN1 in females may be axonal, and in AD it appears to be both axonal and further, a redistribution of PIN1.

**Figure 6.**
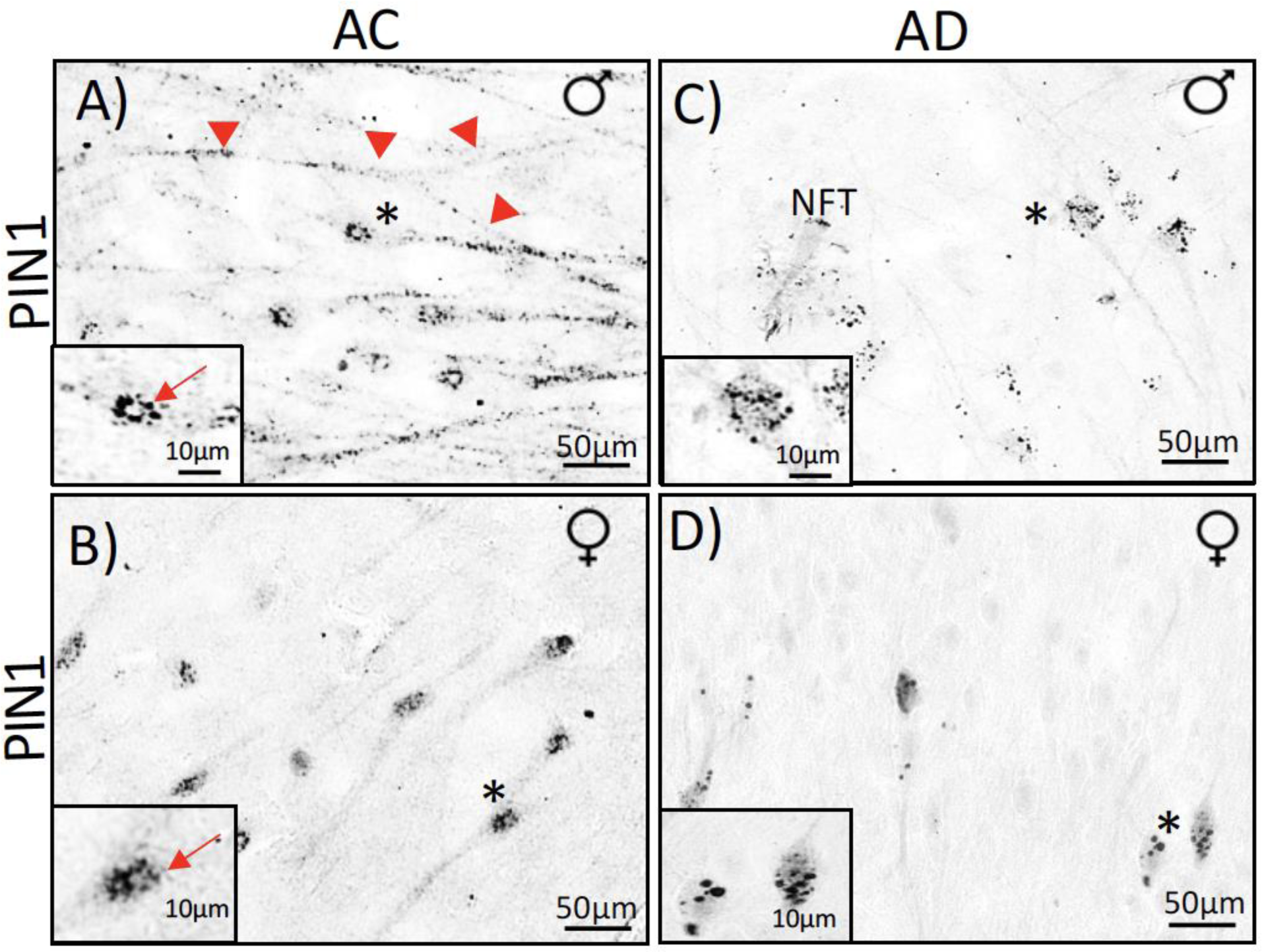
Axonal PIN1 protein levels are decreased in females. Representative photomicrographs of PIN1 immunoreactivity (IR) in temporal neocortex in AD and age matched control (AC) male (♂) and female (♀) samples (A-D). Both AD and AC samples showed positive IR for PIN1. Normal distribution of PIN1 in control samples were clumps of Nissl substance indicative of rough ER (red arrow, A & B insert). Punctate IR in AC extends into the axons (red arrowheads, A). In AC females, this axonal extension of PIN1 is reduced compared to AC males (A vs. B). In AD, PIN1 IR appears to be redistributed in the cytosolic fraction in both male and female AD (C & D insert). AD males show low axonal IR compared to AC males (A vs. C), but female AD subjects are largely devoid of axonal IR (D).

### 3.7. PIN1 colocalizes with the early tau epitope pThr231

The phosphorylation of tau follows a specific pattern of reactivity, accumulating primarily in the entorhinal region in early Braak stages and subsequently progresses to the limbic system (e.g., HIPP), and neocortical regions (e.g., SFG) as disease progresses [40]. The present study focused on the early tau marker pThr231 because of its association with early regional changes [41]. A comparative colocalization study was performed using two highly cited tau antibodies: the late-stage antibody PS396 (Fig.7A) vs. pThr231 an early-stage antibody (**Fig. 7B**) and PIN1 **(Fig. 7C&D)**. The late tau marker (p-tau 396) shows a relatively weak association with PIN1 (6.9% overlap) **Fig. 7E**, and the early tau marker (p-tau 231) shows a very strong association (88.2% overlap) **Fig. 7F**. Interestingly, the cytosolic punctate IR in AD does not colocalize with tangles (**Fig. 7E** insert). It may be that the redistribution of PIN1 may predate the accumulation of phosphorylated tau. There were no significant differences between the number of overlapping cells and sex.

**Figure 7.**
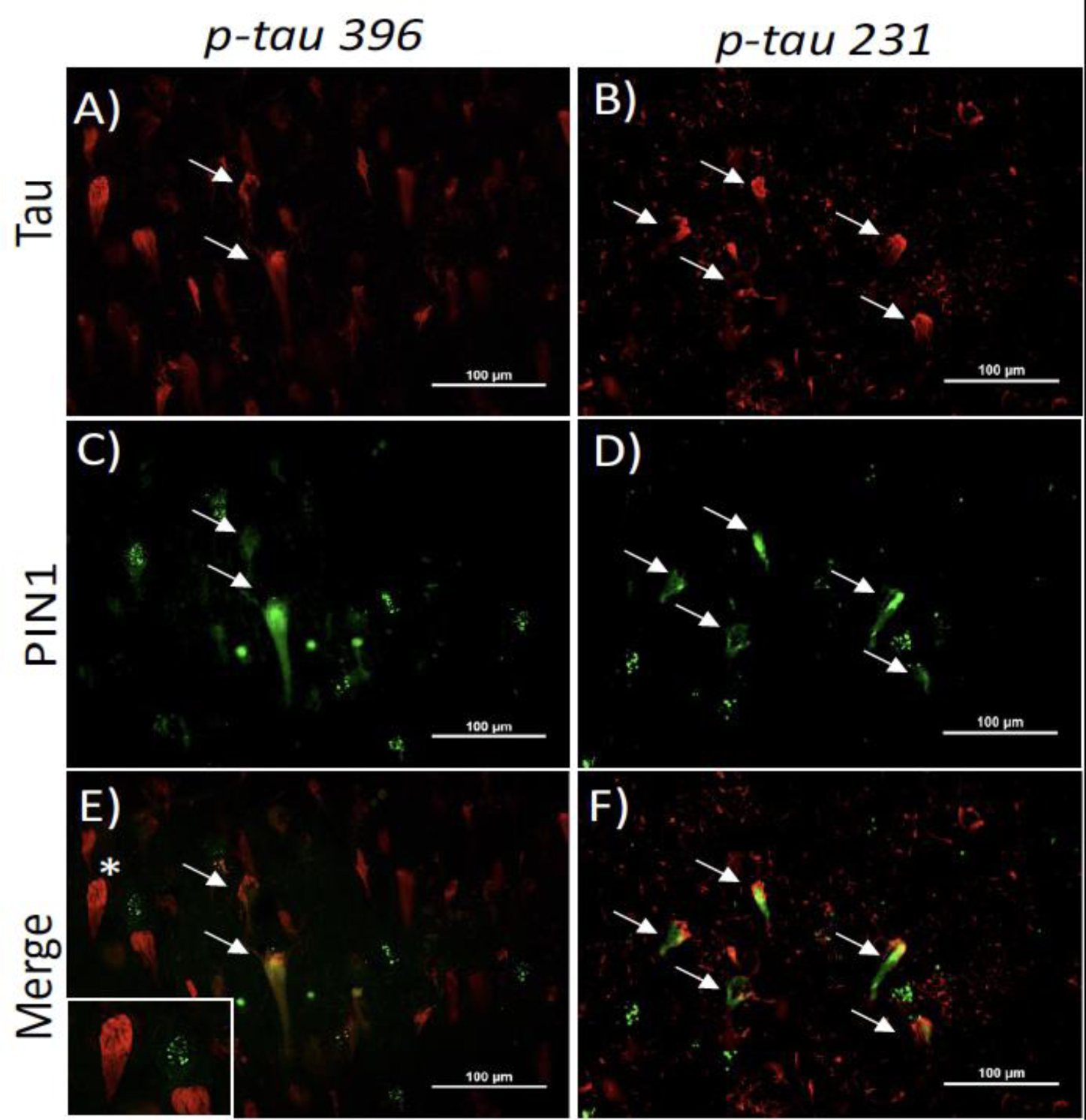
PIN1 colocalizes with the early tau epitope pThr231. Colocalization studies in AD tissue using late tau marker (p-tau 396, (A)) and early tau marker (p-tau 231, (B)) and PIN1 (C & D). The strongest association is with early tau (p-tau 231) (E vs. F). Punctate IR in AD does not colocalize with tangles (E, insert).

3.8. Genetic variation in *PIN1* is nominally associated with risk for early-onset Alzheimer’s disease.

A recent genome sequencing cohort for early-onset Alzheimer’s disease [42] was used to determine if *PIN1* exhibited any genetic association with AD. Analysis was conducted on 227 cases and 671 controls of European ancestry. Rare genetic variation in *PIN1* was nominally enriched in 227 early-onset Alzheimer’s disease cases vs. 671 controls (SKAT p = 0.018, OR [95% CI] = 8.9 [0.7–470.4]). Three of these variants (two in cases, one in a control) were non-coding, but predicted to be damaging by absence from population databases, a CADD score > 10, and presence in a predicted regulatory region (**Supplementary Table 2**). Only one variant was coding but was of particular interest as a predicted stop-gain. All cases were female, consistent with our existing observations.

## 4. DISCUSSION

Imaging [43, 44], cognitive assessment [45, 46], recent peripheral evidence [47], and neuropathological data [48, 49] have established that Alzheimer’s disease affects the brain for decades before clinical diagnosis [50]. Synaptic degeneration and aberrant phosphorylation are some of the earliest pathological features, and strongest correlates to cognitive decline in AD [51, 52, 53, 54, 55, 56, 57, 58, 59]. Although the precise etiology of synaptic degeneration and aberrant phosphorylation in AD are still unknown, recent evidence suggests that *PIN1* may be involved in these processes [11, 12]. Here, we demonstrate that *PIN1* is significantly down-regulated as a function of age (22-99 years) and further downregulated in AD limbic and neocortical brain regions in females more than in males. Additional analysis using the ROS/MAP dataset (N=638 subjects) revealed a decrease in the expression of *PIN1* in the transition to MCI, and a further decrease in the transition to AD, again, driven predominantly by female subjects. We also observed that while *PIN1* expression is negatively correlated with multiregional β-amyloid in both males and females, it is also negatively correlated with multiregional NFT density and global cognitive function only in females. These results suggest the potential utility of therapeutic strategies to augment *PIN1* function in females with preclinical AD and raise the possibility that monitoring *PIN1* could represent an informative disease progression biomarker in female subjects with MCI or AD.

### *PIN1* is a marker of disease progression

Over the past decade, researchers have described the importance of *Pin1* in regulating phosphorylation events associated with AD pathogenesis (reviewed in [10]). Studies of AD animal models and human brain show that Pin1 co-localizes with phosphorylated tau[14, 39, 58] and shows an inverse relationship to the expression of tau [60]. *Pin1* is the only gene, to date, that when deleted can cause both tau and Aβ-related pathologies in an age-dependent manner, in mice [9, 13]. Human studies have shown that allelic distributions of *PIN1* single nucleotide polymorphisms (SNPs), in MCI subjects, represent a risk of developing AD. These findings indicate that polymorphisms of the *PIN1* gene can predict the path to neurodegeneration [61], but it is not clear yet if sex is a contributing factor.

Moreover, AD affects the brain decades before the clinical presentation [49]; our study indicates reduced mRNA *PIN1* levels in amnestic MCI subjects. Whether or not this is a cause or consequence of AD or general aging is unknown. However, it is known that significant *PIN1* compensatory events, such as mitochondrial changes, are taking place in the transition state from MCI to disease [25]. This idea of disease-associated compensation is not a new phenomenon; similar effects were observed in mitochondrial transcripts [25], as well as in another study in early AD [62].

### PIN1, tau, and neurofibrillary tangles

Excessive post-translational modifications (PTMs) (e.g., phosphorylation) are known early events in AD, and studies have shown that some phosphorylation events may be sex-specific [63]. Although phosphorylation events are critical intermediates of “normal” protein function, excessive phosphorylation is pathogenic. Hyperphosphorylation of tau and amyloid triggers the formation of NFTs and toxic Aβ aggregates, both classical hallmarks of AD neuropathology. Recent efforts have identified Pin1 as a key regulator of the phosphorylation signaling pathway in tau [11].

Pin1 regulates the dephosphorylation of tau and APP and promotes microtubule assembly by restoring pThr231-tau ability to bind to them (reviewed in [64]). In distal axons, specifically, Pin1 stabilizes <u>C</u>ollapsin <u>R</u>esponse <u>M</u>ediator <u>P</u>rotein 2A. Importantly, CRMP2A plays a role in translating upstream signaling cascades into axon growth [65]. Therefore, the downregulation of *PIN1* may directly/indirectly affect microtubule assembly and axonal growth, and we provide evidence that this process is sex-specific. We found differences in the distribution of PIN1 protein levels, specifically in axons; AC females showed lower expression, compared to AC males.

Interestingly, PIN1 was compactly wrapped around the nuclear compartment reminiscent of rough ER, in AC males. In AD, PIN1*-*IR appears to be redistributed outside the ER in the cytosolic fraction (in a similar manner in males and females), like what we have observed in other AD studies in our laboratory [28, 29].

Finally, an interesting observation in our laser capture NFT data was that the lowest *PIN1* expression levels were in non-tangle bearing neurons, indicating, but not formally proving that alterations in *PIN1* expression may precede the formation of full-blown NFTs. Further investigations will be required to reconcile our observation of a sex-specific *PIN1* expression difference at the level of tissue homogenate, but an absence at the level of neuronal subtypes (tangle-bearing and non-tangle-bearing). One possibility may be a higher proportion of non-tangle bearing neurons in AD females than AD males, although this remains to be clarified.

Our colocalization studies also show the strongest association with p-tau 231 compared to the ghost tangle marker ps396, the latest tau marker (reviewed in [66]). In a recent study by Ashton and colleagues, the p-tau 231 assay identifies the clinical stages of AD and neuropathology equally well as the earliest reported p-tau marker, p-tau 181. However, the changes in p-tau 231 are increased earlier, before the threshold for amyloid-β PET positivity, and in response to early brain tau deposition [67]. Although these data do not prove that *PIN1* may solely be responsible for the initiation, it is clear there is an association, and it would be well to follow up in a larger cohort, especially in females. Future studies focused on the more toxic oligomeric species of tau could also be warranted. There is evidence that tangles may be a compensatory mechanism to aggregate smaller, more toxic oligomeric species into less toxic inert forms [68].

### *PIN1* expression changes are driven by female subjects

Examination of multiple datasets, multiple brain regions, and multiple classes of cells in multiple disease states clearly shows that females express significantly lower levels of *PIN1* as a function of age and AD. Data from 638 DLPFC samples from the ROS/MAP cohorts demonstrate heterogeneous associations between *PIN1* expression and diverse clinicopathological traits associated with AD, including sex-specific associations. Consistent with our initial analysis, we observed a decline in *PIN1* expression in AD, and subsequent stratification by sex revealed that this decline was driven entirely by female subjects. While we did observe an inverse correlation of *PIN1* with β-Amyloid plaque density and mean SNARE protein density in both males and females, correlations with multiregional tangle density and global cognitive function appear to be female-specific. Given the established literature linking *PIN1* activity with synaptic function and NFT burden in AD generally, these findings may suggest an informative sexual dimorphism in *PIN1* expression in aging females and AD, including impacts on molecular networks that mediate the impact of NFT burden on cognitive function.

Although these findings do suggest that *PIN1* may be involved in some of the earliest events in AD, the exact mechanism by which *PIN1* is dysregulated requires further study. In addition, future research is needed to better understand whether Pin1 ER staining patterns are affected by ER stress or not. Many factors contribute to the expression of genes, from transcription factors to epigenetic modifications (e.g., DNA methylation, microRNAs, histone modifications) to genetic variation. Although rare genetic variation in *PIN1* was nominally enriched in 227 early-onset Alzheimer’s disease cases, further analysis of additional datasets would be beneficial. While further studies are warranted to understand the mechanism of PIN1-mediated regulation of tau phosphorylation (e.g., structural studies to determine the crystal structure of PIN1 in complex with tau and its phosphorylated forms), these results support the potential utility of therapeutic strategies to increase the function of *PIN1* in preclinical AD and raise the possibility that monitoring *PIN1* could be used for tracking the progression of healthy aging to MCI to AD (**Fig. 8**).

**Figure 8.**
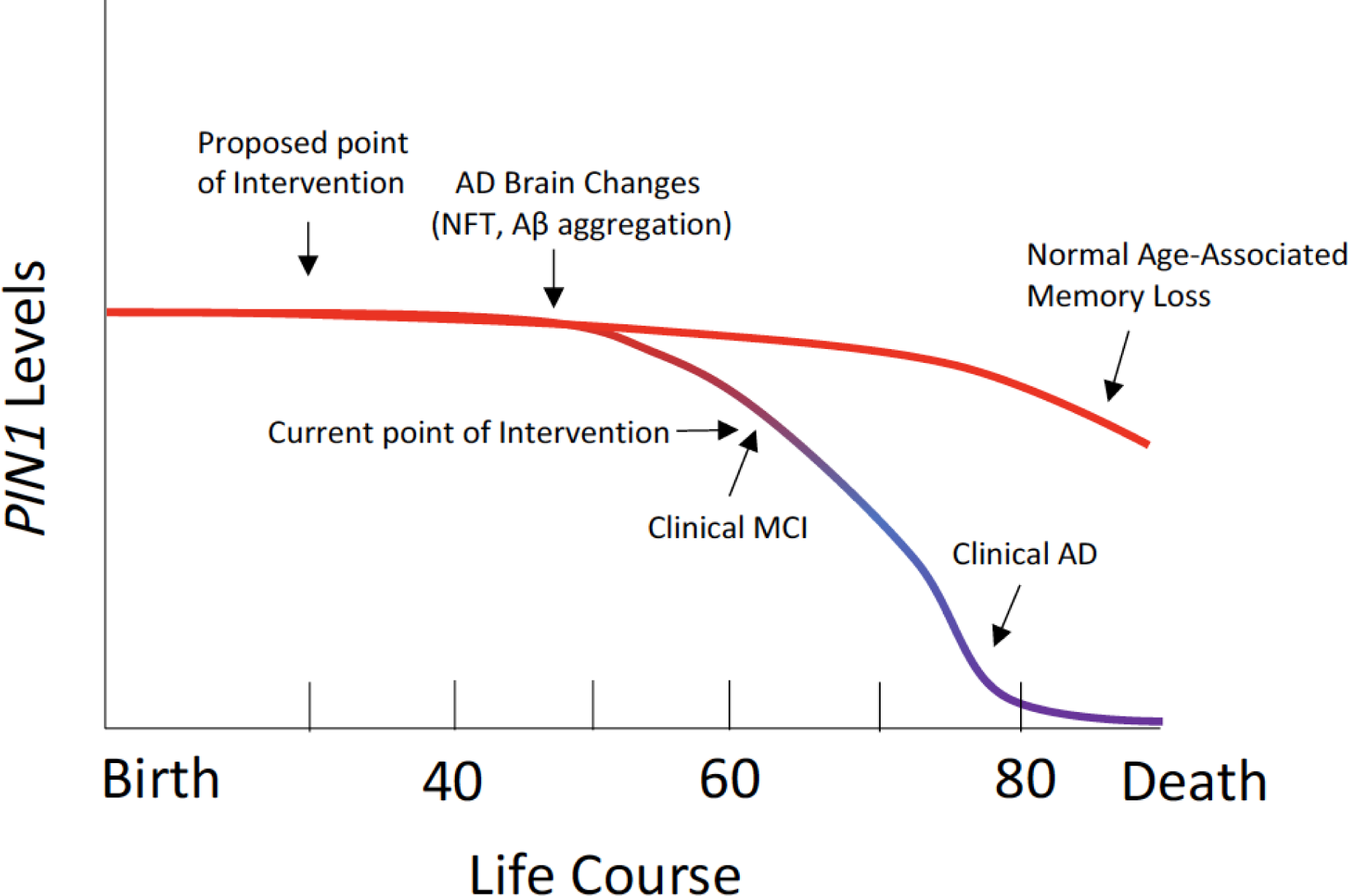
Hypothesized disease progression.

## 5. CONCLUSION

The results show that *PIN1* expression changes occur before clinical symptoms of AD, and correlate to early events associated with AD risk (e.g., synaptic dysfunction), these changes are specific to neurons, and may be a potential prognostic marker to assess AD risk in the aging population and even more so in AD females with increased risk of AD.

## List of abbreviations

Aβ: Amyloid-β peptide
AC: Aged control
AD: Alzheimer’s disease
AMP-AD: Accelerating medicines partnership - Alzheimer’s disease
APP: Amyloid precursor protein’s variants
CRMP2A: Collapsin response mediator protein 2A
DAB: 3.3’- Diaminobenzidine
DLPFC: dorsolateral prefrontal cortex
EC: Entorhinal cortex
ER: endoplasmic reticulum
HIPP: Hippocampus
IR: Immunoreactivity
KO: Knockout
MAP: Memory and aging project
MCI: Mild cognitive impairment
ND: Non-demented
NFTs: Neurofibrillary tangles
ON: Overnight
PBS: Phosphate buffered saline
PIN1: Peptidyl-prolyl cis/transisomerase
PTMs: Post-translational modifications
ROS: Religious orders study
RT: Room temperature
SAPE: Streptavidin phycoerythrin
SFG: Superior frontal gyrus
SNAP-25: Synaptosomal-associated protein-25
SNARE: Soluble N-ethylmaleimide-sensitive factor attachment protein receptor
SNPs: Single nucleotide polymorphisms
VAMP: Vesicle-associated membrane protein
YC: Young control

## Ethics approval and consent to participate

Written informed consent for autopsy was obtained in compliance with institutional guidelines of Banner Sun Health Research Institute. Banner Sun Health Research Institute Review Board approved this study including recruitment, enrollment, and autopsy procedures. Individual person(s) and their respective next-of-kin consented to brain autopsy for the purpose of research analysis as participants in the Banner Sun Health Research Institute brain and body donation program. The human brain tissue used in this manuscript was from routine existing autopsies, which fully qualifies for 4C exemption by NIH guidelines. In addition, samples were analyzed anonymously (e.g. sample numbers) throughout the experimental process.

## Consent for publication

Not applicable

## Availability of data and materials

The results published here are in part based on data obtained from the AD Knowledge Portal (https://adknowledgeportal.org). ROS/MAP study data were provided by the Rush Alzheimer’s Disease Center, Rush University Medical Center, Chicago. Additional phenotypic data can be requested at www.radc.rush.edu. Genome data used in this study is available via the Synapse AD Knowledge portal via accession number syn25686500 and in sources outlined in reference [42].

## Authors’ contributions

CA, wrote manuscript, manuscript preparation for publication, and statistics; CS, performed experiments, data analysis; JN, performed experiments, data analysis JNC: bioinformatics, data analysis; QW, bioinformatics; RV, manuscript preparation/review, data analysis; ED, manuscript preparation/review, data analysis; BH, bioinformatics, data analysis, manuscript preparation/review; DM, conceived of the work, wrote manuscript.

## Acknowledgments

The authors declare no competing financial or conflict of interests. We are grateful to the Banner Sun Health Research Institute Brain and Body Donation Program of Sun City, Arizona for the provision of human biological materials.

