## Supplemental data for "Reduced *PIN1* gene expression in neocortical and limbic brain regions in female Alzheimer’s patients correlates with cognitive and neuropathological phenotypes"

| IHC/Western Sample Metadata (24 AD vs. 24 ND) | | | | |
|---|---|---|---|---|
| | AD Females | AD Males | ND Females | ND Males |
| Sample Size (n) | 12 | 12 | 12 | 12 |
| Expired Age (years) | 80.25 ± 3.1 | 77.5 ± 2.24 | 81 ± 15.62 | 74.7 ± 2.08 |
| Post-mortem Interval (hours) | 3.34 ± .36 | 3.06 ± 0.51 | 2.44 ± .19 | 2.63 ± .462 |
| ApoE Genotype | 2/3 = 0<br>3/3 =10<br>3/4 = 2<br>4/4 = 0 | 2/3 = 0<br>3/3 = 10<br>3/4 = 2<br>4/4 = 0 | 2/3 = 1<br>3/3 = 11<br>3/4 = 0<br>4/4 = 0 | 2/3 = 2<br>3/3 = 10<br>3/4 = 0<br>4/4 = 0 |
| Braak Stage | I = 0<br>II = 0<br>III = 0<br>IV = 2<br>V = 9<br>VI = 1 | I = 0<br>II = 0<br>III = 0<br>IV = 2<br>V = 9<br>VI =1 | I = 1<br>II = 2<br>III = 9<br>IV = 0<br>V = 0<br>VI = 0 | I = 0<br>II = 3<br>III = 9<br>IV = 0<br>V = 0<br>VI = 0 |
| MMSE (last test score) | 6.50 ± 8.74 | 6.80 ± 4.56 | 29.81 ± 0.28 | 28.37 ± 1.2 |

**Supplementary Table 1. Metadata**

| Chr. | hg19 Pos. | Ref. | Alt. | Location / Consequence | CADD | EOAD/Ctrl | Sex |
|---|---|---|---|---|---|---|---|
| 19 | 9949114 | C | T | Exon 2, NP_006212.1 p.(Arg21*) | 37.0 | EOAD | F |
| 19 | 9953695 | G | A | Intron 2 (Deep Intronic) | 14.7 | EOAD | F |
| 19 | 9960702 | G | A | 3' Downstream | 10.1 | EOAD | F |
| 19 | 9961039 | C | CTCCCATCT GGCTGGCT | 3' Downstream | 11.3 | Control | M |

**Supplementary Table 2. Genetic variation in *PIN1* is nominally associated with risk for early-onset Alzheimer's disease.**

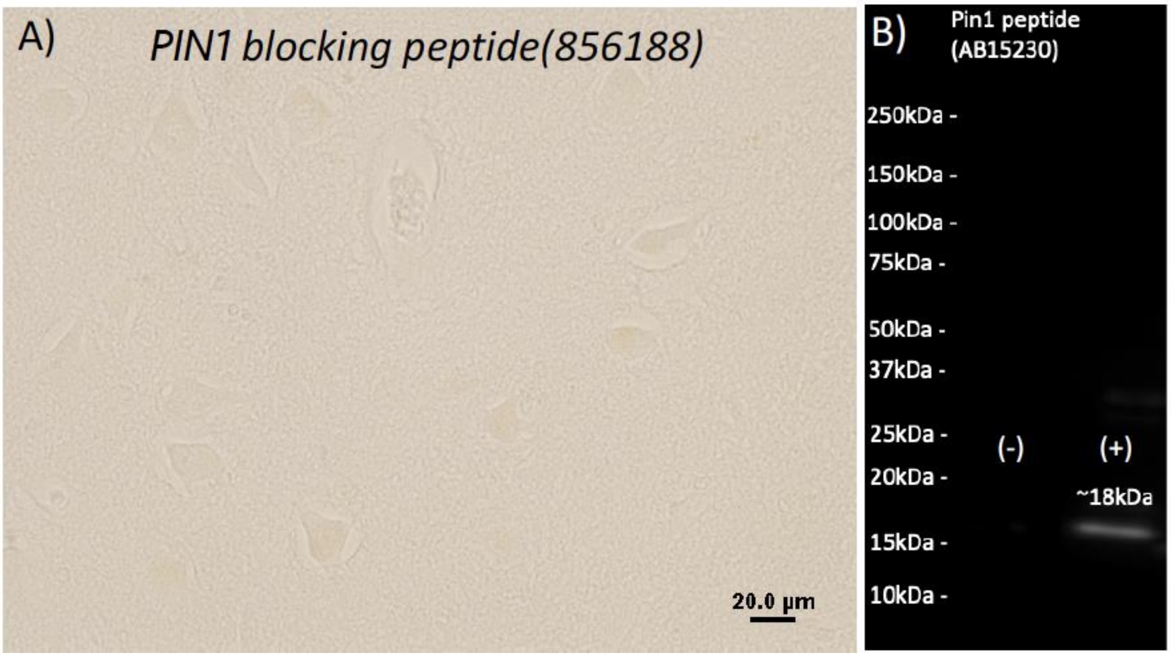


**Supplementary Figure 1. Confirmation of PIN1 antibody specificity**. Specificity was confirmed using PIN1 blocking Peptide (856188) in tissue (A) and in western blot analysis (B). For western blot -/+ = (-). PIN1 Peptide incubated with blocking peptide before running and (+) = PIN1 peptide neat.
